# Carbon footprint of South Dakota dairy production system and assessment of mitigation options

**DOI:** 10.1101/2022.05.16.492173

**Authors:** Anna M. Naranjo, Heidi Sieverding, David Clay, Ermias Kebreab

## Abstract

Livestock production contributes to greenhouse gas (GHG) emissions. However, there is a considerable variability in the carbon footprint associated with livestock production. Site specific estimates of GHG emissions are needed to accurately focus GHG emission reduction efforts. A holistic approach must be taken to assess the full environmental impact of livestock production using appropriate geographical scale. The objective of this study was to determine baseline GHG emissions from dairy production in South Dakota using a life cycle assessment (LCA) approach. A cradle-to-farm gate LCA was used to estimate the GHG emissions to produce 1 kg of energy-and-protein corrected milk (ECM) in South Dakota. The system boundary was divided into feed production, farm management, enteric methane emissions, and manure management as these activities are the main contributors to the overall GHG emissions. The production of 1 kg ECM in South Dakota dairies was estimated to emit 1.21 kg CO_2_ equivalents. The major contributors were enteric methane emissions (46.3%) and manure management (32.6%). Feed production and farm management made up 13.9 and 7.2 %, respectively. The estimate was similar to the national average but slightly higher than the California dairy system. The source of corn used in the dairies influences the footprint. For example, South Dakota corn had fewer GHG emissions than grain produced in Iowa. Therefore, locally and more sustainably sourced feed input will contribute to further reducing the environmental impacts. Improvements in efficiency of milk production through better genetics, nutrition animal welfare and feed production are expected to further reduce the carbon footprint of South Dakota dairies. Furthermore, use of feed additives and anaerobic digesters will reduce emissions from enteric and manure sources, respectively.

## Introduction

Animal agriculture production systems provide food, fiber, and other products to people and can protect and restore carbon (C) in pasture and rangelands while providing wildlife habitat and maintaining other ecosystems [1]. Milk production is the third largest agricultural industry in the U.S., one that has seen tremendous improvement in efficiency over the last century [2]. South Dakota produced 141.8 kt of milk in 2020, which is about 1.4% of total US milk production [3]. Milk production in South Dakota rose by 12% from December 2019 to December 2020, and farmers added about 14,000 new dairy cows during that one-year period continuing a decade-long trend of dairy expansion [3]. These gains came through successful State-level subsidized loan programs and technical assistance to support agri-business investments in modernization as well as improved genetics, nutrition, and animal management.

The dairy industry has an impact on the environment through production of greenhouse gases (GHG) and reactive nitrogen while utilizing natural resources such as land, water, and fossil fuel. For example, livestock contributed an estimated 3.9% of U.S. GHG emissions in 2020 mostly from enteric fermentation, manure storage and field application, and feed production [4]. Several studies have reported mitigation options to reduce the environmental impact of dairy production [5]. Over the last few decades, increased animal productivity reduced the environmental impact of the dairy industry when calculated on an emission intensity (GHG emission per unit of product) basis. For example, Naranjo et al. [6], reported that production of 1 kg of energy- and protein-corrected milk (ECM) in California emitted 1.12 to 1.16 kg of CO_2_ equivalents (CO_2_e) in 2014 compared with 2.11 kg of CO_2_e in 1964, a reduction of 45.0 to 46.9% over the last 50 years. Methane emissions can also be directly reduced through dietary manipulation, feed additives or genomic selection while manure GHG emissions can be reduced through the use of alternative manure management storage techniques and the use of anaerobic digesters [7].

The objective of this study was to determine baseline GHG emissions from dairy production in South Dakota using a life cycle assessment (LCA) approach and assess mitigation options that may reduce the impact of the dairy industry on the environment.

## Materials and methods

An LCA was conducted according to ISO 14040 and 14044 standards [8,9]. The Livestock Environmental Assessment Partnership (LEAP) guidelines by the Food and Agricultural Organization of the United Nations (FAO) [10] were followed for comparability and uniformity with other studies. In accordance with the LEAP guidelines, the Intergovernmental Panel for Climate Change (IPCC) recommendations for reporting global warming potentials [11,12] were used. The IPCC Assessment Report 5 characterization factors were used to report the global warming potentials [13].

### System boundary

The system analyzed in this study covers South Dakota milk production system. The base year for the model was 2018. The South Dakota dairy LCA model included four categories within its system boundary: (i) feed production, (ii) farm management, (iii) enteric methane, and (iv) manure management (Fig 1). All upstream processes such as feed production inputs, as well as extraction and production of fuels were considered in this model up to the farm-gate. Activities post farm-gate, such as milk processing, co-product production, distribution and retail were not included in the model as practices vary widely.

**Fig 1.**
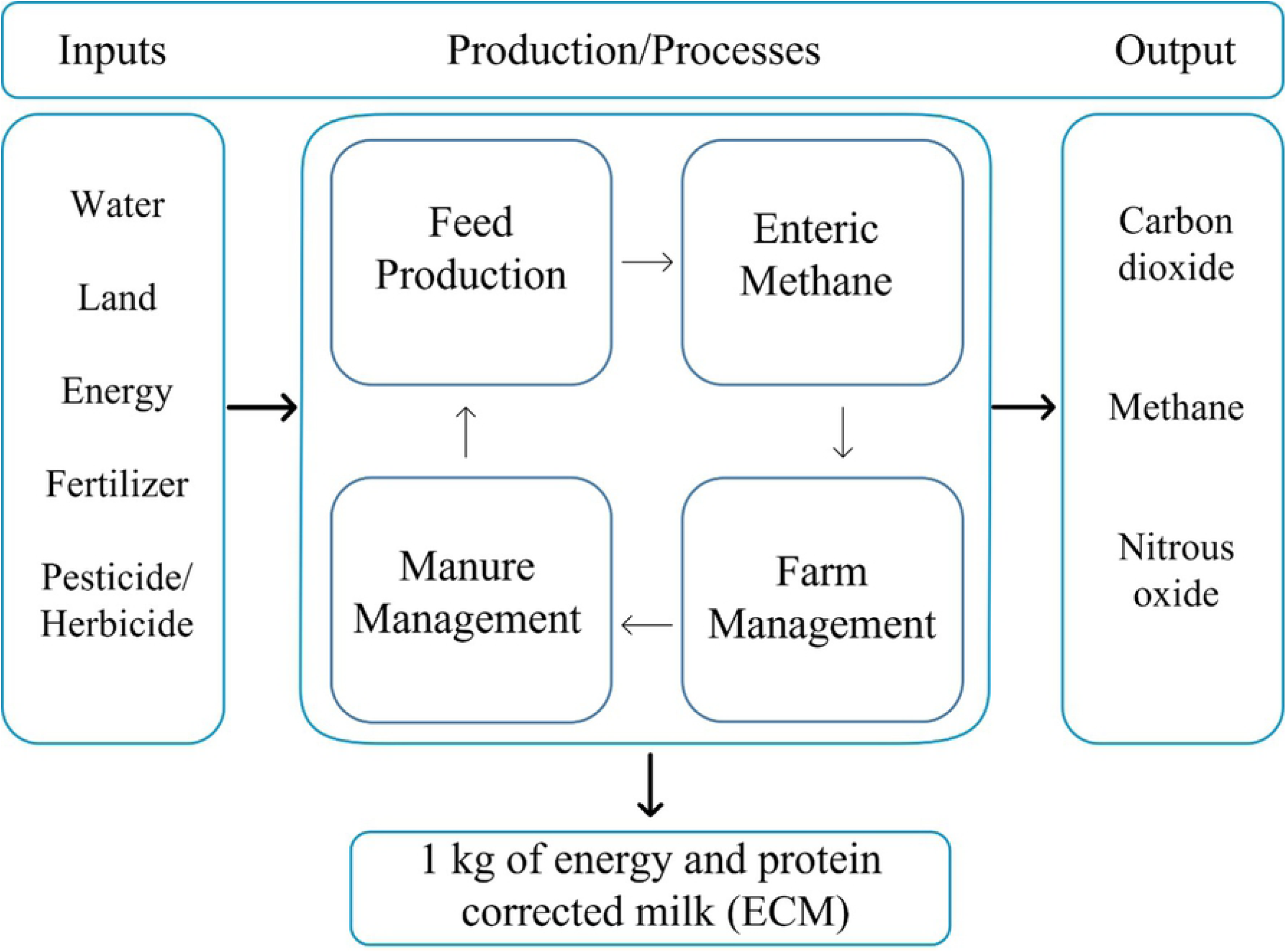
Overview of the milk production system boundary considered in this study.

### Functional unit

The functional unit was 1 kilogram of ECM at the farm gate to standardize the amount of milk production due to the varying nutrient composition of milk from different farms. The ECM was calculated by multiplying the amount of milk production by the ratio of the energy content of the milk to the energy content of standard milk, which is 4% fat and 3.3% true protein [5] as follows:

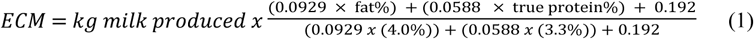

where fat% and protein% are fat and protein percentages in milk, respectively. All the processes in the system were calculated based on a kilogram of ECM. The values for milk production were retrieved form the NASS Quick stats [14] report on average milk production per cow per year in South Dakota from 2016 to 2019. The average annual milk production per cow was reported as 10,131 kg with a composition of 4% fat and 3.2% protein [15]. The standardized ECM for this LCA model was 32.7 kg ECM/day.

### Data sources

The data for this model were collected from multiple sources including the USDA National Agricultural Statistical Service [14], peer-reviewed literature and South Dakota State University (SDSU) extension publications related to diet composition, SDSU dairy extension agents for localized dietary adaptations and common farm practices, Natural Resources Conservation Service for South Dakota (NRCS), Environmental Protection Agency energy emission values [16], and the California Air Resources Board anaerobic digester protocol [17]. The CO_2_ equivalent emissions were calculated by multiplying the emissions of CO_2_, CH_4_ and N_2_O by their 100-year global warming potential (GWP_100_), based on IPCC assessment report 5 [13] which were 1, 28 and 265, respectively.

### Diet composition

Feed rations that made up the diet composition were collected from published data and from dairy farms in South Dakota. Diets collected for calves, heifers, pregnant heifers, lactating cows and dry cows are given in Table 1. The calves’ diets 7 to 16 weeks were taken from Senevirathne et al. [18]. Diets for calves 16 weeks to 14 months was from Schossow [19]. The pregnant heifer diet from 14 months to ∼2 years (25 months) up until close up heifer was according to Anderson et al. [20]. The close-up cow and dry cow rations were based on Diaz-Royon et al. [21]. The lactating cow diet was collected from Ranathunga et al. [22]. The amount of daily feed at each stage was multiplied by the number of days in the respective stage and then added to determine the total lifetime diet. It is assumed that the total lifetime for South Dakota dairy cows is 1,764 days, averaging 2.5 lactations [23].

**Table 1.** Ingredient and nutrient composition of 1 kg of diet for calves, heifers, close-up heifers, lactating cows, and dry cows in the South Dakota dairy system.

| Item<br>Ingredient % DM | Calves |  | Heifer |  | Cow |  | Weighted<br>Average |
| --- | --- | --- | --- | --- | --- | --- | --- |
|  | 7 -16<br>weeks | 4 -14<br>months | Pregnant | Close<br>up | Lactating | Dry |  |
| Alfalfa Hay |  |  |  | 14.2 | 19.1 |  | 14.5 |
| Grass hay |  |  | 40.4 |  |  | 25.2 | 5.47 |
| Beet pulp |  |  |  | 12.3 |  |  | 0.64 |
| Corn grains | 35.0 | 27.6 | 16.2 | 7.92 | 17.1 |  | 16.9 |
| Corn silage |  |  | 25.3 | 42.7 | 38.2 | 42.0 | 34.0 |
| DDGS |  |  |  |  | 5.48 | 3.36 | 4.06 |
| Whole cottonseed |  |  |  |  | 7.86 |  | 5.67 |
| Molasses | 5.10 | 3.49 |  |  |  |  | 0.29 |
| Soybean hulls |  | 6.39 |  |  |  |  | 0.47 |
| Soybean meal | 23.5 | 21.6 | 18.2 | 8.53 | 12.3 | 9.66 | 13.5 |
| Wheat middlings | 35.7 | 41.0 |  |  |  | 2.94 | 3.32 |
| Wheat straw |  |  |  | 14.2 |  | 16.8 | 1.26 |
| Composition |  |  |  |  |  |  |  |
| DMI (kg/d) | 2.00 | 6.00 | 9.00 | 11.0 | 27.1 | 13.2 | 13.4 |
| Days in pen | 67 | 299 | 305 | 123 | 630 | 62 | 1485 |
| DM% | 88 | 88.5 | 63.0 | 20.2 | 49.0 | 53.8 | 57.2 |
| NDF (% DM) | 20 | 24.7 | 39.5 | 43.0 | 35 | 49.5 | 34.5 |
| ADF (% DM) | 7.9 | 10.9 | 24.4 | 22.7 | 23.5 | 31.4 | 19.4 |
| Ether Extract (% DM) | 3.5 | 3.5 | 3.40 | 2.5 | 4.6 | 2.9 | 3.72 |
| CP (% DM) | 21.9 | 22 | 17.8 | 14.1 | 18.7 | 14.2 | 18.2 |

### Crop production

The crop production category includes all processes required to produce the crop and transport it to the farm. Processes include the production of seeds, fertilizers, pesticides or herbicides, energy for machinery, transport, irrigation etc. Transportation is included for instate and out of state crop production. The emissions for transport were determined by doubling the distance traveled depending on crop and the emission factor of that transport method. Tables 2 and 3 provide the region of production, mode of transportation, distance traveled and the yield for crops and byproduct feeds acquired from SDSU Extension calculators and a conversation with dairy field specialist Tracey Erickson, MS (12^th^ June 2020). Data used for crop yields and proportion of production per area were obtained from USDA NASS [14]. Emission factors for crop production inputs such as seed, fertilizers, fuel for machinery were taken from Lal [24]. This study did not include C sequestration in the model, as there is debate over the length of time the C is stored, and models that do include the sequestration of C do not account for the release of it later.

**Table 2.**
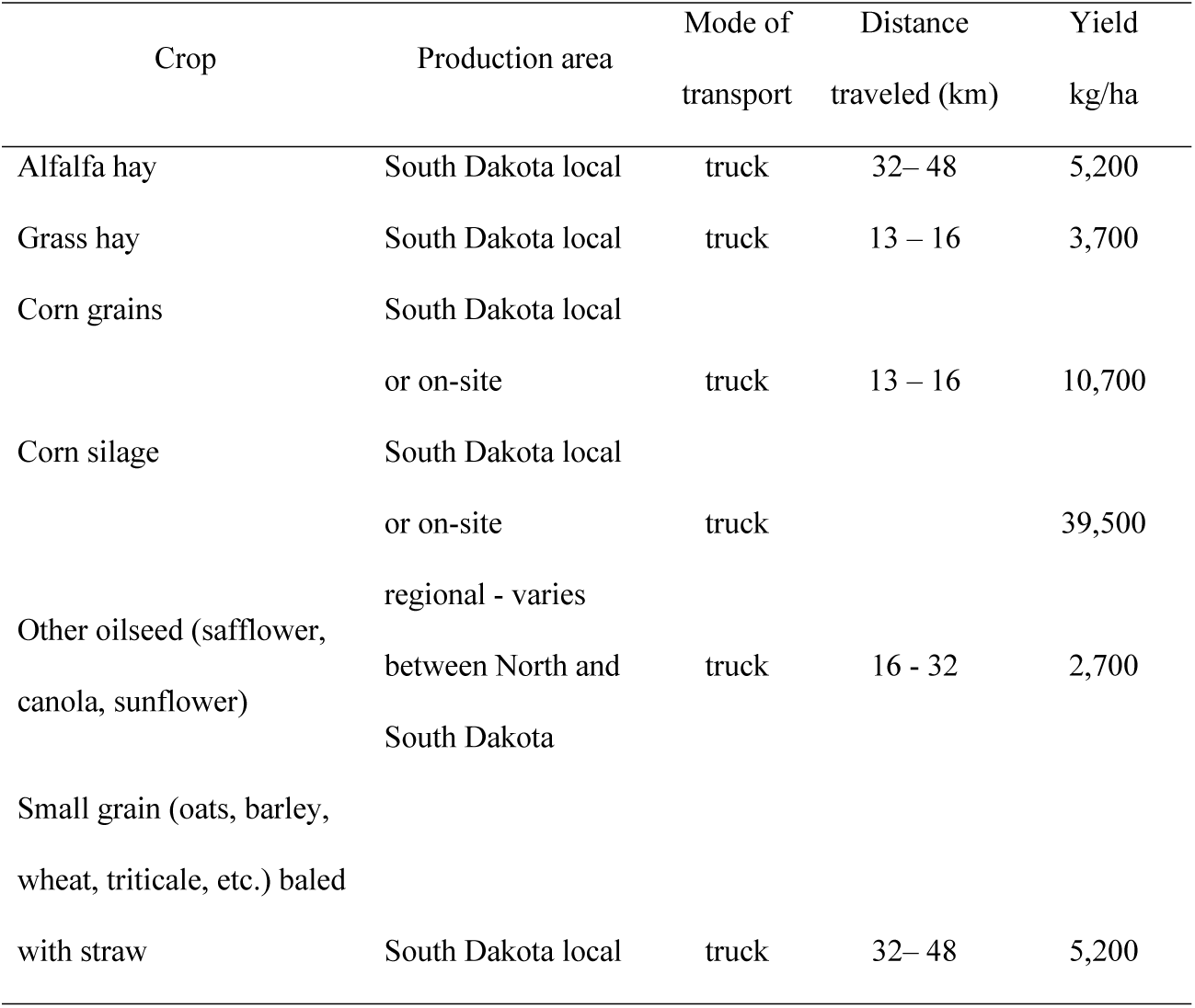
Crop production area, transportation mode, distance traveled and yield.

**Table 3.**
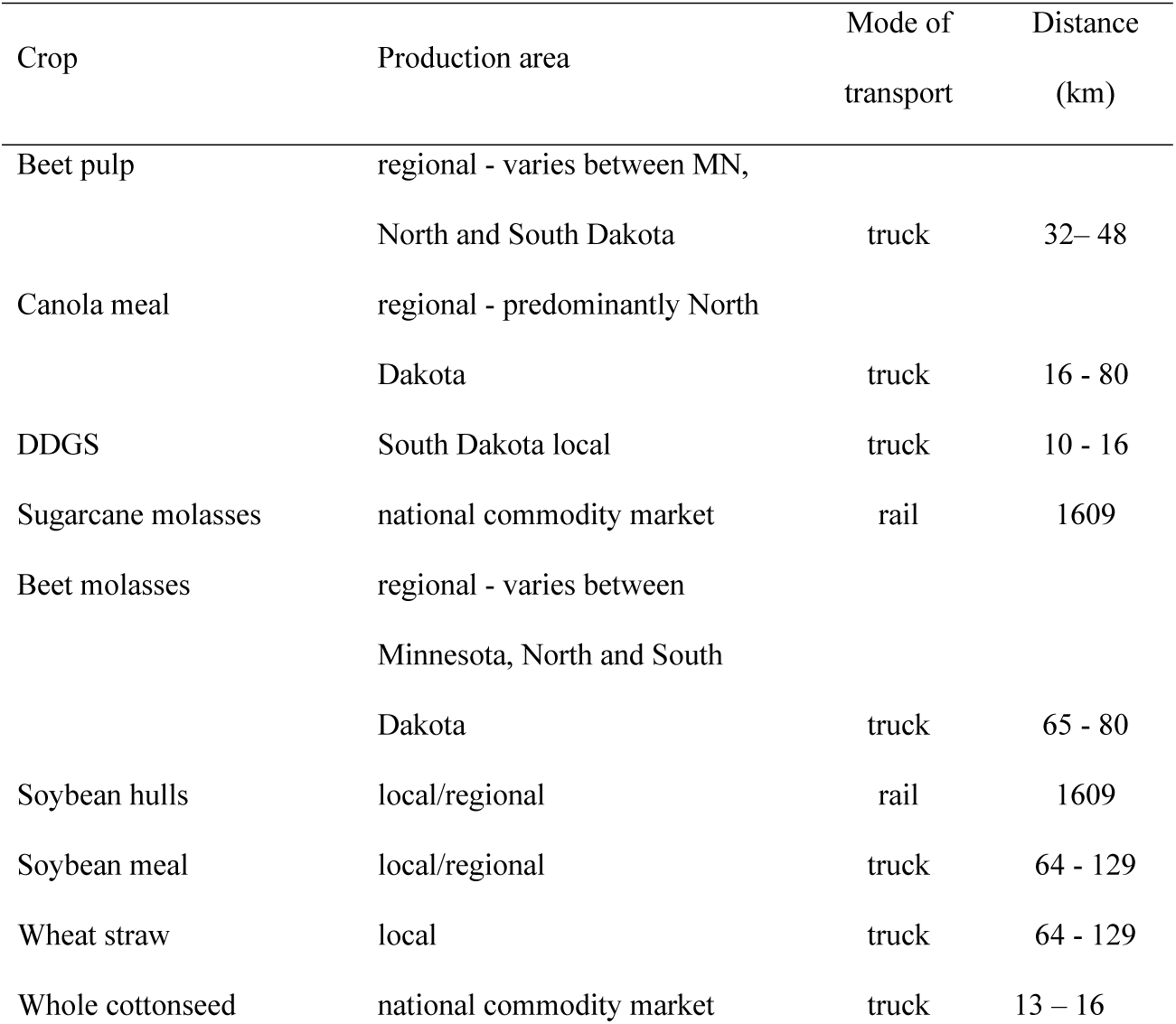
Coproduct or byproduct production region, mode of transportation and distance traveled.

| Crop | Production area | Mode of transport | Distance (km) |
| --- | --- | --- | --- |
| Beet pulp | regional - varies between MN, North and South Dakota | truck | 32– 48 |
| Canola meal | regional - predominantly North Dakota | truck | 16 - 80 |
| DDGS | South Dakota local | truck | 10 - 16 |
| Sugarcane molasses | national commodity market | rail | 1609 |
| Beet molasses | regional - varies between Minnesota, North and South Dakota | truck | 65 - 80 |
| Soybean hulls | local/regional | rail | 1609 |
| Soybean meal | local/regional | truck | 64 - 129 |
| Wheat straw | local | truck | 64 - 129 |
| Whole cottonseed | national commodity market | truck | 13 – 16 |

### Farm management

Farm management includes activities that use energy and water on farm. Energy and water use on-farm are used for animal comfort, milk cooling, cleaning facilities, and collection and transport of manure and waste. Water consumption was estimated according to equations developed by Appuhamy et al. [25]. Different equations were used to estimate water intake from calves after a conversation with J.A.D.R.N. Appuhamy, PhD (November 2016). All three equations are given in Table 4. The free water intake equations required data on sodium and potassium concentrations of the diet. Due to lack of mineral premix composition data, default values were used based on data collected from 43 California dairies [26].

**Table 4.** Equations used to calculate water intake by calves, heifers, dry cows and lactating cows.

| Stage of life | Equation for free water intake (L/d) | Source |
| --- | --- | --- |
| Calves | $2.28 + 0.569 \text{ Age} + 9.58 \text{ Milk Replacer Presence} + 0.037 \text{ DM of Milk Replacer} + 0.0184 \text{ Age}^2$ | Pers. Comm. J.A.D.R.N.<br>Appuhamy, PhD<br>(November 2016) |
| Heifers and dry cows | $1.16 \text{ DMI} + 0.23 \text{ DM}\% + 0.44 \text{ TMP} + 0.061 \text{ TMPC}^2$ | Appuhamy et al. [26] |
| Lactating cows | $-91.1 + 2.93 \text{ DMI} + 0.61 \text{ DM}\% + 0.062 \text{ NaK} + 2.49 \text{ CP} + 0.76 \text{ TMP}$ | Appuhamy et al. [26] |

<sup>2</sup>NaK = joint concentration of Na and K in the diet (mEq/kg of DM), CP = crude protein content (% of DM), NI= Nitrogen intake (g/d), TMP = daily mean ambient temperature, TMPC2 = daily mean ambient temperature squared.

Water (non-free water intake) and energy use on farm was estimated from a Midwest Dairy production guidebook [27]. Energy emission values were collected from the EPA Greenhouse Gases Equivalencies Calculator, based on the eGRID [28]. Farm water and fuel usage reports do not differentiate between water and energy used for milking or other on-farm practices. Therefore, the water and energy use cannot be allocated to other activities.

The South Dakota electricity generation fuel mix falls under the MROW subregion. Compared to the national fuel mix South Dakota uses more coal at 43.9%, less natural gas at 11.2, as well as more wind at 25.1% [28, 29]. South Dakota electricity consumption from 2018 to 2021 showed a decline in use of coal from 47.4% in 2018 to 39.6% in 2020 and an increase in wind power usage from 15.1% to 20.1%, respectively. The emission factor used for electricity is from the EPA’s eGRID summary tables for 2019 South Dakota specific fuel mix [28]. The above average use of coal in the electricity mix in South Dakota contributes to a higher emission factor for electricity than the national average.

### Enteric methane

Enteric methane was estimated using the Appuhamy et al. [30] equations, where the authors evaluated 40 empirical models using region specific data. The best performing model for North America was a modified equation from Nielson et al. [31]. However, due to lack of data on digestible NDF in our dataset, the original unchanged Nielson et al. [31] model was used to estimate enteric methane emissions in South Dakota. The enteric methane (g/d) was estimated using the following equation:

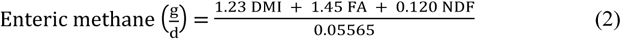

where DMI is dry matter intake (kg/d), FA is dietary fatty acid content (% dry matter) and NDF is dietary neutral detergent fiber (% dry matter).

### Manure management

The methodology for calculating emissions from manure management were based on the IPCC [11] guidelines, with the exception of two equations. The IPCC recommendation for estimating methane from manure requires an estimate of volatile solids (VS) using gross energy intake. Appuhamy et al. [32] developed a model based on North American data, which performed better than the IPCC equation at estimating VS. Therefore, the Appuhamy et al. [32] equation for VS was used to calculate manure methane emissions.

The methane producing capacity (B_0_) for manure from dairy cows and heifers was 0.24 m^3^/kg of VS. The methane conversion factors (MCF) are an estimate of the manure C (in energy terms) that can be converted to methane dependent on the type of storage and climate. The average temperature in South Dakota high dairy producing counties is a four-year average of 45 degrees Fahrenheit or 7.2 degrees Celsius [34], which corresponds to the IPCC 2019 climate zone of *cool temperate and dry*. Manure management systems proportions were updated based on personal communication with USDA Natural Resources Conservation Service (NRCS) South Dakota specialists Justin Bonnema and John Lentz (March 1, 2021). Table 5 shows the proportion of manure management systems reported by the EPA [34] and the NRCS survey on manure management. Additionally, the IPCC [11] updated methane conversion factors for a cool dry climate are shown. Moreover, natural crust covers over anaerobic lagoons can promote methane oxidation by converting methane to carbon dioxide, reducing the methane emissions from the lagoon [35]. Crust covers can form as the top layer is exposed to the air or can be created with the addition of materials such as straw bedding [35]. A notable difference used in this model, was to consider that the winter months in South Dakota create a frozen crust cover over anaerobic lagoons, which acts like a cover, effecting the MCF of anaerobic lagoons. A 40% reduction can be applied due to a crust over when thick and dry [11]. Given the intensity of the winters in South Dakota, the anaerobic lagoons can be completely frozen acting as a crust cover. Therefore, the MCF for anaerobic lagoons was reduced by 40% during the 6-winter months.

The IPCC 2019 refinement guidelines were also followed to calculate direct and indirect nitrous oxide (N_2_O) emissions. A region-specific equation for total nitrogen excretion [36] was used to estimate output instead of the using the IPCC nitrogen excretion equations. Johnson et al. [36] evaluated 45 nitrogen excretion models and determined that an equation by Reed et al. [37] for nitrogen excretion performed best based on multiple datasets. So the equation by Reed et al. [36] was used in this study to estimate nitrogen excretion, which is an input to calculate direct and indirect nitrous oxide emissions using the IPCC [11] equations.

**Table 5.** The proportion of manure management system reported by EPA and NRCS in South Dakota. Methane conversion factor (MCF) from IPCC (2019) is also provided.

|  | EPA | NRCS | MCF |
| --- | --- | --- | --- |
| Lactating and Dry |  |  |  |
| cows | % | % | % |
| anaerobic digester | 0 | 0 | 1 |
| anaerobic lagoon | 54 | 65 | 46.9* |
| daily spread | 0 | 2 | 0.1 |
| deep pit | 14 | 1 | 26 |
| liquid slurry | 2 | 2 | 26 |
| pasture | 14 | 5 | 0.47 |
| solid storage | 16 | 15 | 2 |
| dry lot | 0 | 10 | 1 |
| Calves and Heifers |  |  |  |
| anaerobic digester | 0 | 0 | 1 |
| anaerobic lagoon | 0 | 0 | 67 |
| daily spread | 8 | 5 | 0.1 |
| deep pit | 0 | 2 | 26 |
| liquid slurry | 0 | 0 | 26 |
| pasture | 5 | 10 | 0.47 |
| solid storage | 0 | 5 | 2 |
| dry lot | 87 | 78 | 1 |

\*The adjusted anaerobic lagoon MCF corresponds to a natural cover which covers the lagoons in South Dakota during the winter months. The default MCF for a cool climate is 66% and emissions are halved with a cover.

### Allocation of Co-product

Beef is a co-product of the dairy industry; therefore, the environmental impacts and resource use were allocated between dairy and beef production systems. The International Dairy Federation [38] recommend biophysical allocation, which splits the burden between milk and meat based on the total amount of milk a cow produces in her lifetime and the live weight of the cow at slaughter. In this model the final weight of cows is assumed to be 580 kg, with allocation of 90% for milk and 10% for meat production. The allocation factor was applied to categories determined to be for both products. In this model the crop production, enteric methane and manure management emissions were allocated using the biophysical factor. However, for the farm management category, the energy and water data were not split based on milking and non-milking activities, so the milk production carries the full burden of that category.

### Assumptions/ limitations

The results of this LCA are limited to the above defined system boundary and scope. The LCA is based on the average animal performance and does not include the animal-to-animal variation. Whenever possible, primary data from surveys and reports on dairy farmers in South Dakota were used, otherwise published literature was used. Additionally, emission factors for South Dakota were used, however, due to the lack of data, some factors were developed from a neighboring state, or close-by regions such as the Midwest or national U.S. values. Readily accessible and centralized emission factors would prove useful for future studies.

Care should be taken when comparing this study with others because this LCA model used the IPCC [13] assessment report 5 values for methane and N_2_O. Additionally, we modified 3 equations (for estimating enteric methane, VS, and total nitrogen excretions) in the IPCC methodology for calculating emissions from livestock. The exclusion of other life cycle impact categories may result in an incomplete snapshot of the overall performance of the products. Social and economic impact categories were not considered in this model, so trade-offs between environmental, social, and economic factors were not evaluated.

## Results and discussion

### Greenhouse gas emissions

The production of 1 kg of ECM in the South Dakota milk production system was associated with 1.22 kg CO_2_e in 2018 (Fig 2). The emissions for milk production in South Dakota were just slightly lower compared to the national US LCA reported by Thoma et al. [39] of 1.23 kg CO_2_e/kg ECM. However, Thoma et al. [39] used lower GWP values for methane emissions (25) instead of the updated value of 28 kg of CO_2_ equivalencies, which may indicate that the South Dakota GHG emissions per kg of ECM may even be lower than the national average. The South Dakota estimate was slightly more than was estimated for California milk production systems, which ranged between 1.12 to 1.16 kg CO_2_e in 2014 [6].

**Fig 2.**
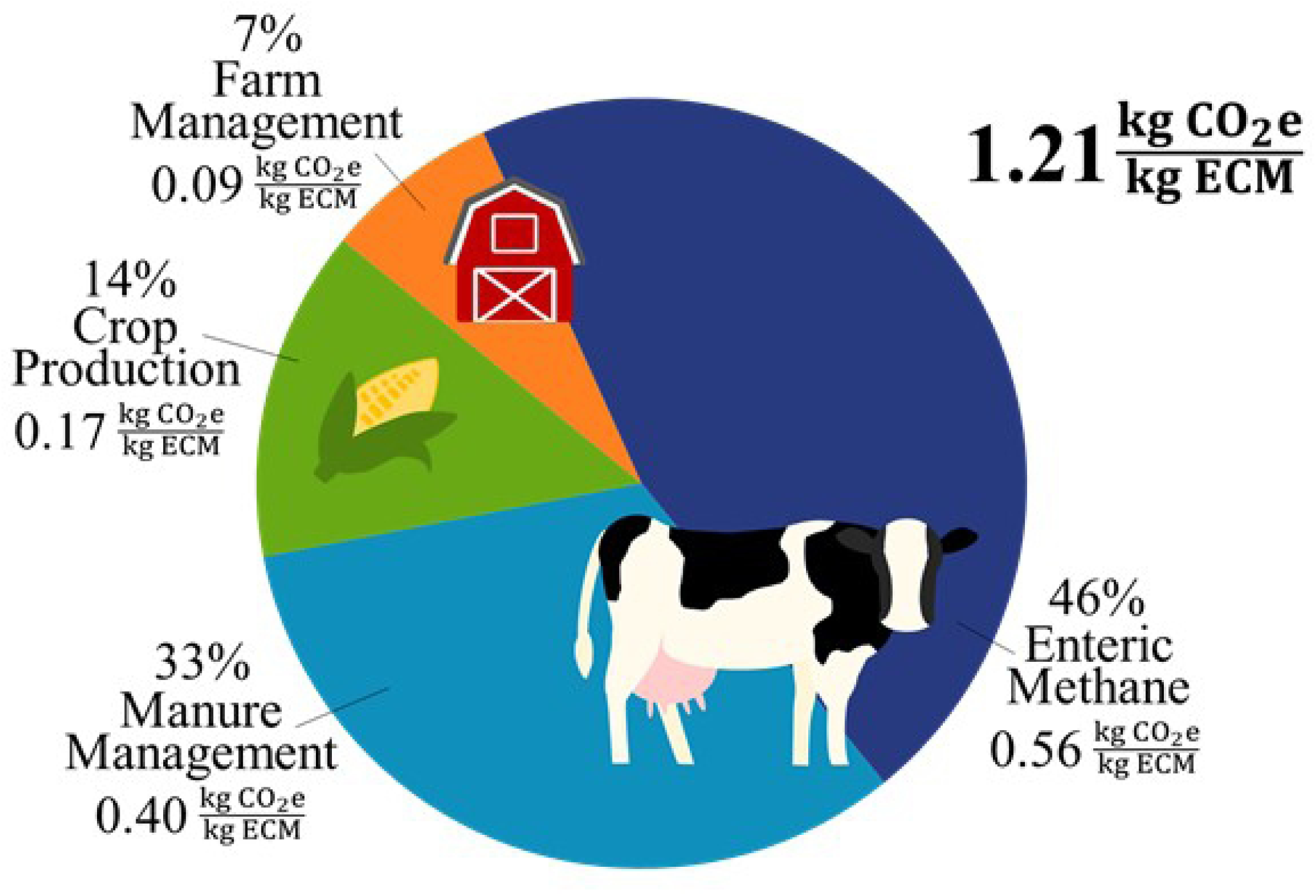
Greenhouse gas emissions using Global Warming Potential_100_ to produce 1kg of energy corrected milk (ECM) in South Dakota.

The largest contributor to emissions in South Dakota was enteric methane making up 46.1% of total GHG emissions. Manure management was the next leading category at 32.9% of total GHG emissions. Crop production (13.8%) and farm management (7.2%) were the third and fourth sources of GHG emissions.

### Emissions from crop production

The emissions from crop production were 13.8% of the total emissions. When compared to Thoma et al. [39] study, the South Dakota contribution to crop production was lower (0.17 kg CO_2_e/kg ECM) than the crop contribution in Region 4 of the US model (0.33 kg CO_2_e/kg ECM). This is mainly due to crop yield improvements since 2008, which was the reference year for Thoma et al. [39] study and composition of diet. The lifetime rations were primarily comprised of corn silage (34.0%), corn grains (16.9%) and alfalfa hay (14.5%) (Table 1). By-product use in South Dakota was relatively low at 5.98% compared to the average byproduct use of 10 states reported by Russomanno et al. [40], which averaged at 31.3% and ranged from 12.7 to 56.7% of total diet. Greater inclusion of byproducts in dairy diets would reduce the C footprint as they do not carry an environmental burden apart from further processing and transportation related emissions.

The feed efficiency of South Dakota cows was lower compared to California which required an average of 1.0 kg of feed to produce 1 kg ECM, while California dairies have a feed-to-milk ratio of up to 0.80 kg feed per kg ECM [6]. A smaller ratio means that it requires less feed to produce one kg ECM. A further reduction in feed-to-milk ratio through either inclusion of more digestible feed or genetic improvement would further reduce the C footprint of South Dakota dairies.

### Corn production

A common ingredient in the South Dakota rations is corn grains. Most corn grains in the US are produced in the Midwest states, particularly, Iowa, Illinois, Minnesota, and Indiana. Corn grains are then transported to farms across the nation from the Midwest for use in dairy cattle rations. Some studies looking at many Midwest corn grain production include C sequestration into the calculations. However, there is a debate between researchers, as many consider this C a temporary storage. Additionally, through sensitivity analyses it has been shown the final emission factor for the grain is influenced greatly by the amount of soil C included in the estimate [41]. Lee et al. [41] shows an emission factor for South Dakota corn grain production to be between −1.72 and 1.31 kg CO_2_e/kg corn, and a greater range (−6.4 to 20.2 kg CO_2_e/kg corn) for all U.S. Midwest production. Due to the large differences in emission factors and the sensitivity to including C sequestration, our model excludes it. Pelton’s corn production study examines almost all U.S. states to assess any differences between corn production at the county level focusing on fertilizer use, N_2_O emissions and irrigation emissions while holding other inputs constant [42]. Although, this study is the next step-in region-specific emission factors, the estimates for corn production are much smaller, for example for South Dakota the estimate is 0.053 kg CO_2_e/kg corn [42].

To further examine the emissions for corn grain production, this study investigated whether corn grain production in South Dakota was more efficient than Iowa corn grain production and if this results in lower emissions. South Dakota corn grain production inputs were from the South Dakota State University Extension Crop Budgets [43]. Due to the limited reporting of crop production inputs such as fertilizer quantity, pesticide use, seed and energy used for machinery, only Iowa corn production reports were used for comparison as they have the most complete information. The Iowa corn grain production inputs were taken from the Iowa State Extension reports on Estimated Costs of Crop Production [44]. The Iowa corn grain production inputs for corn were averaged over four years (2017-2020) and used to calculate the corn grain emission factor.

Fig 3 shows the comparison between South Dakota and Iowa corn grain production. The biggest difference in production is that Iowa corn grains had greater fertilizer use, which also affects emissions from managed soils. The emission factors for corn grain production in South Dakota and Iowa are 0.28 and 0.32 kg CO_2_e/kg crop, respectively. If corn grains from Iowa were to be used instead of South Dakota corn grain, the estimated emissions from crop production would increase from 0.169 to 0.174 kg CO_2_e/kg crop. However, the increase in crop production related emissions only increases the total GHG emissions by 0.41%.

**Fig 3.**
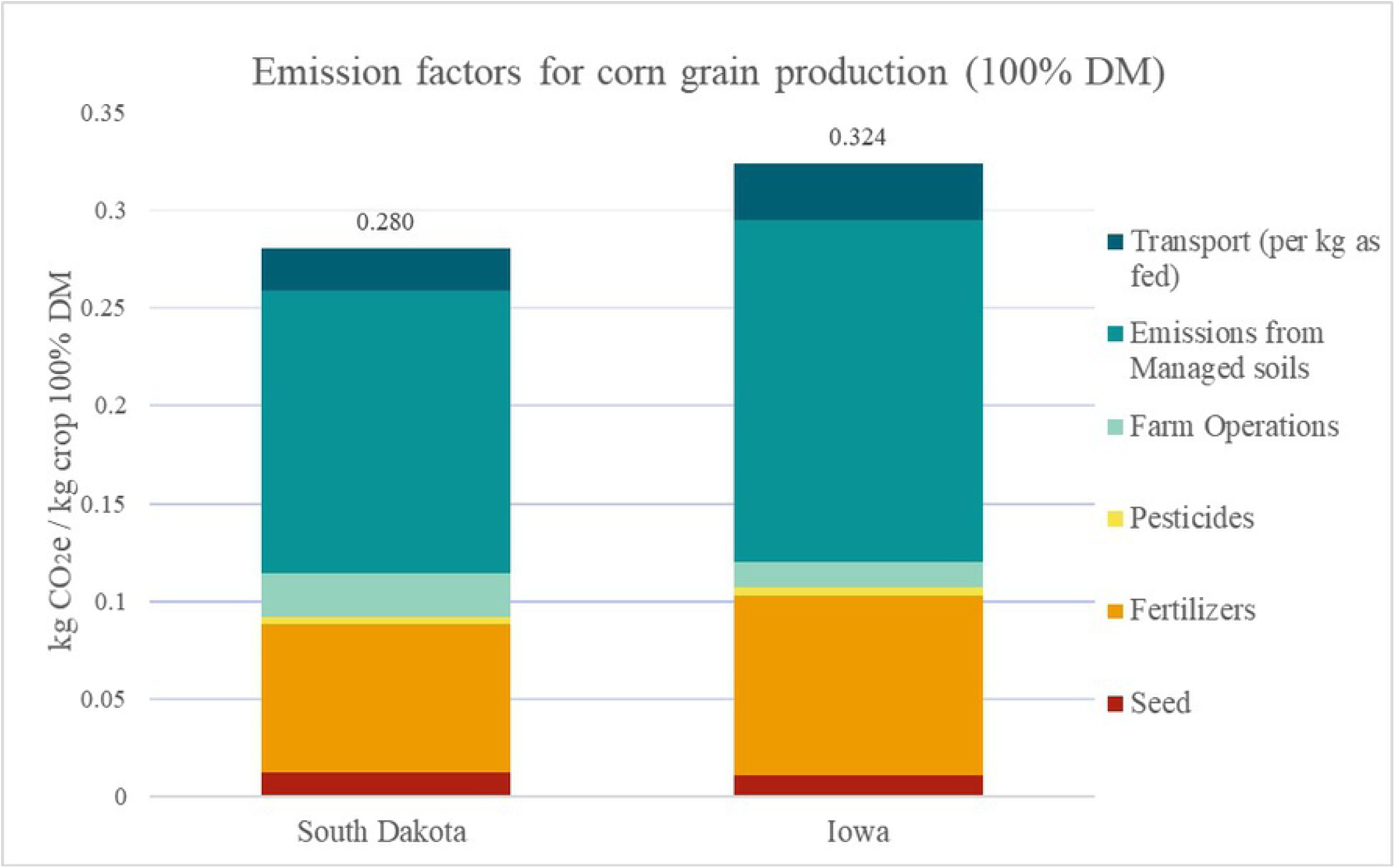
Comparison of emissions related to corn grain production at the farm gate in South Dakota and Iowa.

### Emissions from farm management

Emissions from farm management are the smallest contributor to overall farm GHG emissions, which was in agreement with previous studies reported [6,39]. This category included the energy, such as electricity and natural gas for required for cooling and storing milk, heating water for cleaning and any other activity on farm that require energy. As more energy is sourced from renewable methods such as solar and wind, whether it’s at the farm level or state level, the emissions from this category will be reduced.

### Emissions from enteric methane

Enteric methane is the largest contributor to total emissions (46.1%). The inclusion of high forage in the ration (58.6%) explains the relatively greater emissions from enteric fermentation. This value is in line with the Thoma et al. [39] model, which found that enteric methane was the largest contributor to total emissions in the US national dairy LCA. In contrast, the California model estimated that enteric methane emission was the 2^nd^ largest contributor to total on-farm emissions. However, in California, most dairies use manure lagoons, and it is relatively warmer throughout the year, which carry a greater MCF value, and hence more methane is emitted in manure storage. The model used to estimate the national enteric methane emissions [39] was an equation that includes an input of total forage proportion in the diet. The current study goes a step further and used and equation that takes into consideration the fiber content of the diet not just forage, which was evaluated to have a better accuracy [30].

There are several mitigation options that can be implemented to reduce enteric methane emissions. For example, Kebreab and Fouts [45] describe that improving forage quality and increasing the proportion of concentrates in the diet leads to lower enteric methane emissions. Honan et al. [46] also reviewed several feed additives that directly reduce enteric methane emissions. Additives such as 3-NOP have been estimated to reduce enteric emissions by over 40% in dairy cattle [47]; while the macroalgae *Asparagopsis spp*. reduced emissions by 67% in dairy cattle [48] and over 80% in beef cattle [49], however none of the feed additives are commercially available at present. A combination of several mitigation options may be used in South Dakota dairy systems and contribute to considerable reduction in enteric methane emissions.

### Emissions from manure management

The emissions from manure management contribute 32.9% to whole farm emissions. Emissions from manure management were dependent on how manure is handled and stored. Solid based manure storage systems have much lower MCF compared to liquid-based storage. The recent NRCS data showed that there was a shift with decreasing solid manure storage and increasing liquid manure storage than previously estimated by EPA (Table 6). Specifically, anaerobic lagoon storage increased from 54% to 65%, which contributed largely to GHG emissions as the MCF of anaerobic lagoons is much greater compared to other forms of storing manure. As warmer weather becomes more common, the winter months may shorten, and in-turn reduce the number of months the anaerobic lagoons are frozen. If a shorter winter five-month occurred, the emissions from manure management would increase from 0.40 to 0.43 kg CO_2_e/kg ECM increasing total emissions to 1.25 kg CO_2_e/kg ECM. Similarly, a four-month winter would increase manure management emissions to 0.46 kg CO_2_e/kg ECM and total emissions to 1.28 kg CO_2_e/kg ECM. Assuming the trend continues in increased use of anaerobic lagoons and temperature, increased emissions from manure management is expected in the future.

Although there are several methods of reducing methane emissions from dairy production systems such as solid-liquid separation and manure additives [7], anaerobic digesters have been gaining popularity due to their potential to reduce methane emissions, generate renewable energy as well as increase the fertilizer value of digestate. In California, the Low Carbon Fuel Standard (LCFS) program has incentivized the development of anaerobic digesters and use of biogas from dairy farms to power vehicles [50]. In the current study, we analyzed the impact of constructing anaerobic digesters in South Dakota.

### Anaerobic digesters

The EPA’s AgSTAR program is proponent of anaerobic digester use for the recovery of biogas and reduction in methane emissions from manure [51]. To explore this as a mitigation strategy, a consequential model was used to determine how the implementations of a digester can impact GHG emissions. We used the anaerobic digester system enterprise budget calculator from the Center for Sustaining Agriculture and Natural Resources at Washington State University [52], which does not include the digester construction. The default anaerobic digester was used but can be modified using manufacturer provided values. The methane output was assumed to be 0.23 m^3^ CH_4_ per kg VS and 3.53 kWh/ m^3^ CH_4_ conversion factor was used. As the proportion of manure for lactating and dry cows increased the anaerobic digester proportion, the anaerobic lagoon proportion decreased proportionally. Additionally, the electricity and heat recovery from combustion of captured methane were used to meet the on-farm energy demands, further decreasing net impacts. Fig 4 shows how manure management and farm management emissions change as anaerobic digester use replaces manure storage in anaerobic lagoon. The baseline model had 0.40 kg CO_2_e/kg ECM from manure management, and at the high value of 65% of anaerobic digester usage, the manure management emissions decrease to 0.03 kg CO_2_e/kg ECM. Additionally, the farm can meet its own energy demands further reducing emissions from farm management. Therefore, the use of anaerobic digesters in South Dakota dairy farms has the potential to reduce total GHG emissions by about 30%.

**Fig 4.**
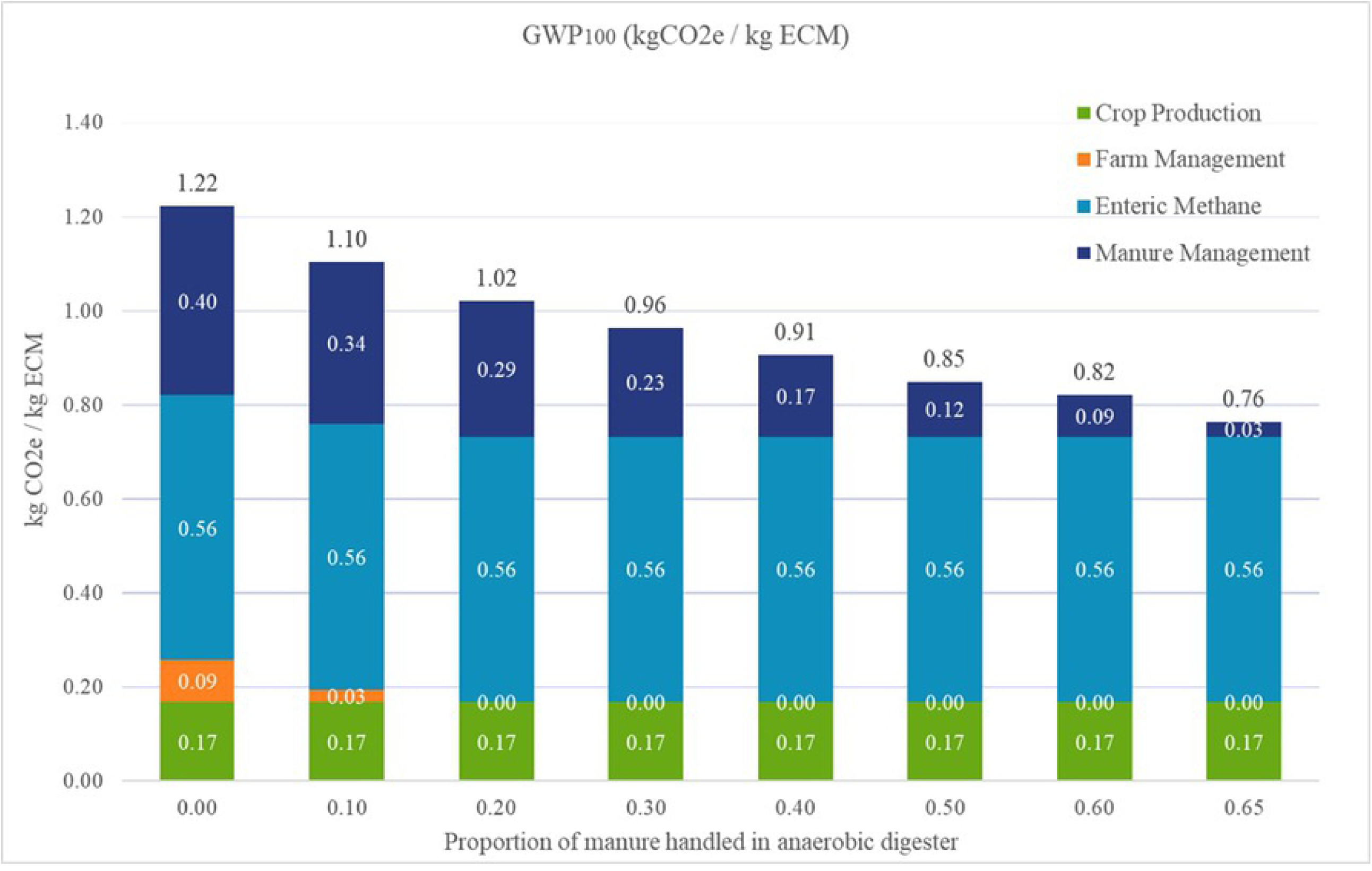
Total emissions per kg ECM as anaerobic digester use increases.

## Acknowledgments

The study was supported by South Dakota Corn Utilization Council.

## Notes

### Competing Interest Statement

The authors have declared no competing interest.

